# The engineered peptide construct NCAM1-Aβ inhibits aggregation of the human prion protein (PrP)

**DOI:** 10.1101/2021.01.04.425177

**Authors:** Maciej Gielnik, Lilia Zhukova, Igor Zhukov, Astrid Gräslund, Maciej Kozak, Sebastian K.T.S. Wärmländer

## Abstract

In prion diseases, the prion protein (PrP) becomes misfolded and forms fibrillar aggregates, which are resistant to proteinase degradation and become responsible for prion infectivity and pathology. So far, no drug or treatment procedures have been approved for prion disease treatment. We have previously shown that engineered cell-penetrating peptide constructs can reduce the amount of prion aggregates in infected cells. The molecular mechanisms underlying this effect are however unknown. Here, we use atomic force microscopy (AFM) imaging to show that the aggregation of the human PrP protein can be inhibited by equimolar amounts of the 25 residues long engineered peptide construct NCAM1-Aβ.

## 1. Introduction

Prion and amyloid diseases are both characterized by aggregation of misfolded proteins or peptides (Jaunmuktane and Brandner, 2019; Miller, 2009; Sengupta and Udgaonkar, 2018; Verma et al., 2015), such as the prion (PrP) protein (Creutzfeldt-Jakob disease), α-synuclein (Parkinson’s disease), and amyloid-β (Aβ) and tau (Alzheimer’s disease). Many of these proteins and peptides may co-aggregate or at least influence each other’s aggregation (Luo et al., 2016, 2017; Ren et al., 2019; Wallin et al., 2018). Factors that modulate the aggregation of one of these proteins, such as small molecules, potential drug compounds, lipids, and metal ions, can often modulate also the aggregation processes of other proteins in this family (Ambadi Thody et al., 2018; Chemerovski-Glikman et al., 2016; Gielnik et al., 2019; Owen et al., 2019; Richman et al., 2013; Robinson and Pinheiro, 2010; Wallin et al., 2017; Wärmländer et al., 2013; Wärmländer et al., 2019; Österlund et al., 2018). This suggests that the underlying mechanisms may be the same in prion and amyloid diseases (Jaunmuktane and Brandner, 2019; Jucker and Walker, 2018; Miller, 2009). Prion aggregates are however particularly infectious, as they spread between cells (Jaunmuktane and Brandner, 2019; Jucker and Walker, 2018), and are not degraded by cellular processes such as proteinase digestion (Jaunmuktane and Brandner, 2019; Löfgren et al., 2008; Söderberg et al., 2014).

The toxic species in amyloid and prion diseases are generally considered to be small toxic oligomeric aggregates (Sengupta and Udgaonkar, 2018; Verma et al., 2015), but so far no drugs or treatments that target such aggregates have been approved against prion diseases (Hyeon et al., 2020; Lee et al., 2019; Mashima et al., 2020). Potential drug molecules may interfere with oligomer formation in various ways: by reducing production of the protein, by inhibiting its aggregation, by diverting the aggregation pathway(s) towards non-toxic forms, or by reducing the lifetime of the toxic forms, for example by promoting rapid aggregation into larger non-toxic aggregates.

We have previously demonstrated anti-prion properties in short peptide constructs (up to 30 residues) with sequences derived from the unprocessed N-termini of mouse and bovine prion proteins: such PrP-derived peptides induced lower amounts of prion aggregates resistant to proteinase K in prion-infected cells (Löfgren et al., 2008; Söderberg et al., 2014).

The PrP-derived peptides consisted of an N-terminal signal peptide segment (different for mouse and bovine PrP), together with a conserved positively charged and hydrophobic hexapeptide (KKRPKP) corresponding to the first six residues of the processed PrP protein. Our earlier studies had shown that peptides with such sequences were able to interact with and penetrate cell membranes (Lundberg et al., 2002; Magzoub et al., 2005; Magzoub et al., 2006; Oglecka et al., 2008). The anti-prion effects of the PrP-derived peptides were lost when the KKRPKP hexapeptide was coupled to various peptides with cell-penetrating properties (Söderberg et al., 2014). The anti-prion effects were however retained when KKRPKP was coupled to the signal sequence of the Neural cell adhesion molecule-1 (i.e., NCAM1_1-19_) (Söderberg et al., 2014).

The mouse PrP_1-28_ segment and the NCAM1_1-19_-KKRPKP construct are both amyloidogenic in themselves, as they form amyloid fibrils by self-aggregation (Mukundan et al., 2017; Pansieri et al., 2019). The NCAM1_1-19_-KKRPKP construct was recently shown to inhibit aggregation of the amyloid-β peptide involved in Alzheimer’s disease (Henning-Knechtel et al., 2020), and to promote *in vitro* aggregation of the amyloid protein S100A9 (Pansieri et al., 2019), which is involved in amyloid-related and other inflammatory processes (Horvath et al., 2018; Wang et al., 2019; Wang et al., 2014). Almost identical results were obtained for a similar amyloidogenic 25 residue-construct, i.e. NCAMh-19-KKLVFF (from here onwards: NCAM1-Aβ) (Pansieri et al., 2019). The KLVFF sequence originates from the hydrophobic core (residues 16-20) of the Aβ peptide: this pentapeptide is known to inhibit aggregation of the full-length Aβ peptide (Tjernberg et al., 1996). In the NCAM1-Aβ construct, an additional lysine residue was added to the KLVFF sequence for increased solubility (Pansieri et al., 2019). The molecular properties of the NCAM1-Aβ sequence and its segments are shown in Table 1, including hydrophobicity values calculated according to the Wimley-White whole residue hydrophobicity scale (Wang et al., 2016; Wimley and White, 1996).

**Table 1.** Primary sequences and molecular properties of the human PrP protein, the NCAM1-Aβ peptide construct, and its parts.

| Protein | Sequence | Isoelectric point (pI) | Molecular weight [g mol<sup>-1</sup>] | Net charge at pH 7 | Theoretical hydrophobicity [kcal mol<sup>-1</sup>] |
| --- | --- | --- | --- | --- | --- |
| **huPrP<sub>23-231</sub>** | UniProt ID: P04156<br>(209 aa) | 9.39 | 22747 | +7 | - |
| **NCAM1<sub>1-19</sub>-K-Aβ<sub>16-20</sub> (NCAM1-Aβ)** | **NH<sub>2</sub>-MLRTKDLIWTL**<br>**FFLGTAVS**<u>**KKLVFF**</u>**-NH<sub>2</sub>** | 11.67 | 2974.7 | +4 | -3.83 |
| **NCAM1<sub>1-19</sub> (NCAM1)** | **NH<sub>2</sub>-MLRTKDLIWTL**<br>**FFLGTAVS-NH<sub>2</sub>** | 11.39 | 2211.7 | +2 | -3.06 |
| **KKLVFF** | **NH<sub>2</sub>-KKLVFF-COOH** | 10.69 | 781 | +2 | -0.77 |

As the NCAM1-Aβ construct inhibits fibrillation of the Aβ peptide (Henning-Knechtel et al., 2020), but promotes (co-)aggregation of the S100A9 protein (Pansieri et al., 2019), it is unclear how the construct may affect the aggregation of the PrP protein (if at all). Here, we use Atomic Force Microscopy (AFM) imaging to investigate if there is a direct effect of the NCAM1-Aβ construct on the *in vitro* aggregation of the human PrP protein. Answering this question might help clarify the mechanisms underlying the previously observed beneficial effects of such peptide constructs on PrP infectivity (Löfgren et al., 2008; Söderberg et al., 2014).

## 2. Materials and Methods

### 2.1 Sample preparation

Human recombinant prion protein (huPrP) was prepared according to a previously published protocol (Morillas et al., 1999; Zahn et al., 1997), albeit with some modifications. The plasmid contained the full-length (23-231)huPrP protein in fusion with an N-terminal HisTag, and the thrombin cleavage site was cloned into the pRSETB vector (Invitrogen, USA). The construct was expressed in E. Coli (BL21- DE3) grown in LB growth medium with 100 μg/mL ampicillin. Expression was induced by isopropyl β-D-galactopyranoside (IPTG) at OD600 = 0.8. Sonication of the lysates was performed in a buffer containing 100 mM Tris at pH 8, 10 mM K2HPO4, 10 mM glutathione (GSH), 6 M GuHCl, and 0.5 mM phenylmethane sulfonyl fluoride (PMSF). The solution was centrifuged and the supernatant loaded to Ni-NTA resin (GE Healthcare) and eluted with buffer E (100 mM Tris at pH 5.8, 10 mM K2HPO4, and 500 mM imidazole). After washing the resin, the protein was purified with two-step dialysis, initially against 10 mM phosphate buffer with 0.1 mM PMSF at pH 5.8, and then against Milli-Q H2O with 0.1 mM PMSF. After thrombin cleavage, the pure huPrP protein (i.e., with the HisTag removed) was concentrated using an Amicon Ultra 0.5 ml centrifugal filter (Merck & Co., USA) with an NMWL cutoff of 3 kDa. The final protein concentration was determined by spectrophotometry using an extinction coefficient of ε_280_ = 57995 M^−1^cm^−1^ (Gasteiger et al., 2005). The quality of the final protein was controlled by mass spectrometry (molecular mass 22747 Da - Table 1).

The NCAM1-Aβ peptide (Table 1) was purchased as a custom order from the PolyPeptide Group (France) in lyophilized form. The peptide was dissolved in Milli-Q water, and its concentration was determined via triplicate UV absorption measurements at 280 nm, using a DS-11 spectrophotometer (DeNovix, USA) and an extinction coefficient of ε_280_ = 5500 M^−1^cm^−1^ (Gasteiger et al., 2005).

### 2.2 Sample incubation

The initial buffer in the huPrP solution was exchanged to ultrapure water by triplicate diafiltration using an Amicon Ultra 0.5 ml centrifugal filter (Merck & Co., USA) with an NMWL cutoff of 3 kDa. Samples of 0.5 μM NCAM1-Aβ, 2.5 μM NCAM1-Aβ, 0.5 μM huPrP, and 0.5 μM NCAM1-Aβ+ 0.5 μM huPrP were then prepared in 10 mM sodium phosphate buffer, pH 7.5, with 100 mM NaCl and 2 M urea. The urea was added as it has previously been shown to promote unfolding of the native PrP structure, which is the first step towards aggregation (Julien et al., 2009; Swietnicki et al., 2000). The samples were incubated for 72 hours at 50 °C with magnetic stirring at 400 rpm. Subsamples were taken out for AFM imaging (below) after 8 and 72 hours, respectively.

### 2.3 Atomic force microscopy (AFM) imaging

Incubated samples (5 μl) were transferred to freshly cleaved mica plates and left to absorb for 1 min, rinsed three times with 300 μl of pure water, and then dried under a gentle flow of nitrogen. AFM imaging was performed on a JPK Nanowizard 4 (Bruker, Germany) AFM unit using Tap150Al-G cantilevers (Ted Pella Inc., USA) in air intermittent contact mode. The scan rate was 0.3 - 0.7 Hz, the scan area size was 5 μm x 5 μm or 10 μm x 10 μm, with 512 x 512 or 1024 x 1024 pixel resolution respectively. The AFM images were analyzed using the Gwyddion 2.54 software (Necas and Klapetek, 2012).

## 3. Results and Discussion

AFM images of the aggregation products present in the samples after 8 hours of incubation are shown in Figs. 1A-D. The sample of 0.5 μM huPrP readily self-aggregated into long fibrils (Fig. 1A) that are approximately 3 - 4 nm thick (judged by their measured height, as width is not accurately represented in AFM images). This is somewhat thinner but still in line with the results of previous studies on PrP fibrils (Terry and Wadsworth, 2019; Vazquez-Fernandez et al., 2017; Yamaguchi and Kuwata, 2018). A few very large aggregate clumps, over 10 nm high, can also be seen (Fig. 1A). For NCAM1-Aβ, the 0.5 μM sample shows small aggregate clumps (Fig 1B). Some of them are relatively large, with heights over 6 nm, and may or may not be early stages of fibrillar aggregates (Luo et al., 2014). The 2.5 μM NCAM1-Aβ sample shows numerous mature fibrils, about 2 - 3 nm high, together with aggregate clumps (Fig. 1C). The more abundant amount of fibrils for 2.5 μM of NCAM1-Aβ confirms earlier results showing that NCAM1-Aβ self-aggregates faster at higher concentrations (Pansieri et al., 2019).

**Figure 1.**
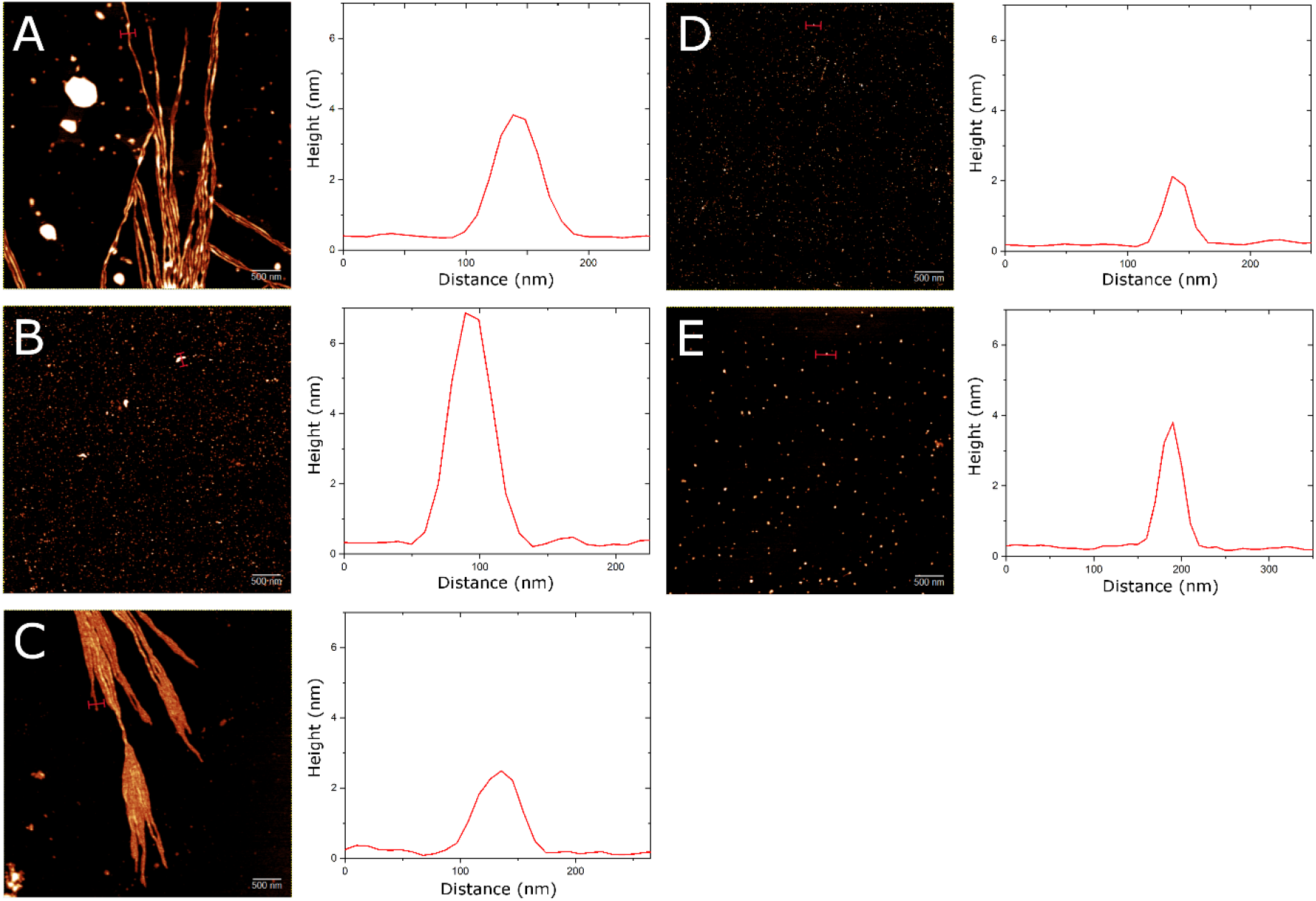
AFM images of: (A) 0.5 μM huPrP protein; (B) 0.5 μM NCAM1-Aβ peptide; (C) 2.5 μM NCAM1-Aβ peptide; and (D) 0.5 μM huPrP protein + 0.5 μM NCAM1-Aβ peptide. All samples in A-D were incubated for 8 hours. (E) 0.5 μM huPrP protein + 0.5 μM NCAM1-Aβ peptide, incubated for 72 hours. All studied samples were incubated at 50 °C in 10 mM sodium phosphate buffer, pH 7.5, with 100 mM NaCl and 2 M urea, and with magnetic stirring at 400 rpm. The white scale bars are 500 nm.

Interestingly, the sample containing both 0.5 μM NCAM1-Aβ and 0.5 μM huPrP displays no fibrils, but only numerous small aggregate clumps, about 2 nm high (Fig. 1D). Even after 72 hours no fibrils can be seen, but the aggregate clumps are then fewer and larger, around 3 - 4 nm high (Fig 1E). As it cannot be ruled out that these small aggregate clumps will eventually form fibrils, it is not possible to tell if fibrillation is completely inhibited, or if the fibrillation rate merely is significantly reduced. Nonetheless, the absence of fibrillar aggregates of huPrP in the presence of equimolar concentrations of NCAM1-Aβ clearly shows that the peptide construct directly interacts with the huPrP protein and interferes with its aggregation. As both molecules are positively charged (Table 1), it stands to reason that they interact mainly via hydrophobic forces.

The aggregation-inhibiting effect of NCAM1-Aβ (Fig. 1) appears to provide an explanation, at a molecular level, to our earlier observations that such peptide constructs significantly reduce the levels of prion aggregates in prion-infected cells (Löfgren et al., 2008; Söderberg et al., 2014). As both the NCAM1-Aβ peptide and the huPrP protein can form amyloid fibrils by themselves (Figs. 1A and 1C), the two molecules may interact via cross-aggregation, to form smaller non-fibrillar co-aggregates (Fig. 1E) that could be less toxic than pure huPrP aggregates (Luo et al., 2016, 2017). If so, the huPrP/NCAM1-Aβ interactions would be similar to the interactions between Aβ and NCAM1-Aβ (Henning-Knechtel et al., 2020). In any case, the huPrP/NCAM1-Aβ interactions are very different from the interactions between NCAM1-Aβ and S100A9 protein, where amyloid aggregation is promoted (Pansieri et al., 2019). Because the NCAM1-Aβ construct has different effects on different aggregating proteins, it would be interesting to study how this construct might affect the aggregation of other disease-related prion proteins, such as those involved in animal diseases like bovine spongiform encephalopathy (BSE), chronic wasting disease in cervids, and sheep scrapie (Vazquez-Fernandez et al., 2017).

## 4. Conclusions

Our atomic force microscopy images show that the *in vitro* aggregation of the human PrP protein is inhibited by equimolar amounts of the 25 residues long engineered peptide NCAM1-Aβ. Thus, a very likely molecular-level explanation to our previous observation that such cell-penetrating peptide constructs can reduce the amount of prion aggregates in infected cells, is that these peptide constructs directly interact with the PrP protein and prevent its fibrillation.

